# Unified comparison of spinal locomotion control architectures in neuromechanical simulations

**DOI:** 10.64898/2026.06.09.731213

**Authors:** Vincent Ton, Seungmoon Song

## Abstract

Neuromechanical simulations provide a powerful framework for investigating how neural control architectures generate and regulate human locomotion. Numerous biologically inspired locomotion controllers have been proposed, including reflex-based, central pattern generator (CPG)-based, and muscle synergy-based models. However, direct comparison across studies remains difficult because of differences in musculoskeletal models, optimization methods, and evaluation protocols. Here, we implemented four representative locomotion control architectures, reflex-based, CPG-reflex-based, muscle synergy-based, and CPG-reflex-synergy-based controllers, within a unified neuromechanical simulation framework to enable controlled comparisons under shared biomechanical and computational conditions. Performance was assessed in terms of (1) agreement with experimentally observed gait characteristics, including kinematics, kinetics, muscle activations, and biomechanical trends across speeds and slopes, and (2) locomotor versatility across speed–slope conditions. The reflex-based and CPG-reflex-synergy-based controllers most closely reproduced experimentally observed gait characteristics, while the CPG-reflex-synergy controller achieved the broadest range of stable walking behaviors across speeds and slopes, followed closely by the reflex-based controller. These findings should be interpreted as comparisons of specific model implementations rather than definitive evaluations of the underlying biological hypotheses. Moreover, because the investigated controllers primarily focused on spinal-level mechanisms for nominal steady-state locomotion, the limited versatility observed in some of the models across broader speed and slope conditions suggests the importance of integrating spinal locomotor mechanisms with supraspinal modulation when modeling locomotion beyond nominal steady gait. To facilitate further investigation, we publicly share the simulation framework and controller implementations.

**Key points:**

- Existing neuromechanical locomotion controllers have been difficult to compare directly because of differences in simulation frameworks.
- We implemented four representative spinal locomotion control models (reflex-based, central pattern generator (CPG)-reflex-based, muscle synergy-based, and CPG-reflex-synergy-based) within a unified simulation framework and compared their human-likeness and versatility.
- The reflex-based and CPG-reflex-synergy-based controllers best reproduced human-like gait characteristics, while the CPG-reflex-synergy-based controller demonstrated the greatest locomotor versatility across speed–slope conditions, followed closely by the reflex-based controller.
- Because the investigated controllers primarily modeled spinal-level mechanisms associated with steady-state locomotion, their reduced adaptability across broader speed and slope conditions highlights the importance of incorporating supraspinal modulation when modeling locomotion beyond nominal gait.
- We publicly share the simulation framework and controller implementations to support further investigation of human locomotion control.

## Introduction

Neuromechanical simulations offer a powerful computational testbed for studying how the nervous system generates and regulates human locomotion, a long-standing scientific problem with far-reaching implications for neurorehabilitation, assistive device design, and human performance enhancement (Song & Geyer, 2018). Experimentally identifying the underlying control algorithms and neural circuitry that produce coordinated walking is not feasible with current techniques. Human locomotion emerges from the interaction between large-scale neural networks and the nonlinear physics of the musculoskeletal system, including muscle tendon dynamics, contact forces, and whole-body mechanics, making it challenging to disentangle the structural organization of control from the resulting movement. Directly measuring or mapping the activity of the vast neuronal populations, particularly during natural unconstrained movement, remains beyond current experimental capability, leaving a substantial gap between measurable neural signals and observable behavior. By coupling physiologically inspired neural control models with physics based musculoskeletal models, neuromechanical simulations provide a complementary approach that enables systematic testing of candidate control architectures and investigation of how specific control principles give rise to locomotor behavior.

A wide range of biologically inspired locomotion control models have been proposed and evaluated in neuromechanical simulations, most of which focus on generating stereotypical steady gait through low-level automatic control mechanisms commonly attributed to spinal circuitry. These mechanisms are typically framed in terms of spinal reflexes, central pattern generators (CPGs), and muscle synergies. Spinal reflex pathways transform proprioceptive and cutaneous sensory signals into rapid, phase-dependent modulation of motor output, and experimental studies in humans using methods such as H-reflex modulation, stretch perturbations, and cutaneous stimulation provide direct evidence for their involvement in regulating gait and responding to perturbations (Dietz, 2002; Nielsen, 2003). CPGs refer to neural circuits capable of generating rhythmic motor patterns through networks of reciprocally interacting neurons, often modeled using coupled neural oscillators; although extensive experimental work in vertebrate animal preparations demonstrates that such circuits can produce locomotor rhythms without sensory feedback, evidence for their functional role in intact human walking remains indirect and debated (Grillner, 2006; Ijspeert, 2008). Muscle synergies, in contrast, are not typically interpreted as mechanisms for generating neural drive, but rather as a strategy for coordinating and distributing motor commands across groups of muscles through low-dimensional activation modules inferred from electromyographic (EMG) recordings (Ivanenko et al., 2004). Circuit-level studies in animal models support the existence of modular spinal organization consistent with muscle synergies, but whether the low-dimensional modules identified from human electromyographic data directly reflect such neural circuitry or instead partly emerge from biomechanical and task constraints remains debated (Tresch & Jarc, 2009). Reflecting these uncertainties, neuromechanical locomotion models often incorporate different combinations of reflex feedback, rhythm-generating circuits, and synergy-based coordination to represent spinal-level locomotor control. Representative implementations of these control principles in neuromechanical simulations are summarized in the subsection *Existing neuromechanical locomotion control models* below.

Although a number of biologically inspired locomotion control models have been developed and investigated, direct comparison across studies remains challenging because methodological differences confound interpretation of their relative performance. Existing models are implemented using diverse musculoskeletal representations that vary in segment definitions and inertial properties, joint configurations, muscle sets, muscle dynamics, musculoskeletal attachment definitions, and contact mechanics, all of which can influence emergent gait behavior. Control parameters are determined using different approaches, ranging from manual or heuristic tuning to numerical optimization. Although optimization provides a more systematic means of parameter selection, studies often employ different cost functions, constraints, and optimization algorithms. Furthermore, simulation protocols, numerical solvers, and evaluation criteria are not uniform across the literature, with some studies focusing on kinematic resemblance, others prioritizing energetic efficiency, and others emphasizing robustness or specific locomotor tasks. Consequently, it remains unclear how different control models compare when evaluated under shared biomechanical and computational conditions.

Here, we implement representative locomotion control models within a unified simulation framework to enable direct and controlled comparisons (Fig. 1). By adopting a common musculoskeletal model, optimization procedure, and evaluation protocol, we assess the simulated gait produced by each control architecture along two key dimensions: human-likeness and versatility. Human-likeness refers to how closely simulated gait reproduces experimentally observed biomechanical and energetic characteristics, whereas versatility reflects the range of speed and slope conditions over which stable gait can be generated. This unified approach allows us to isolate how differences in implemented control architectures within each hypothesis influence gait realism and adaptability under consistent physiological and computational constraints. To facilitate reproducibility and further investigation, we provide our MATLAB/Simulink implementation, including all four control models, through a public repository: *https://doi.org/10.5281/zenodo.20509551*.

**Figure 1:**
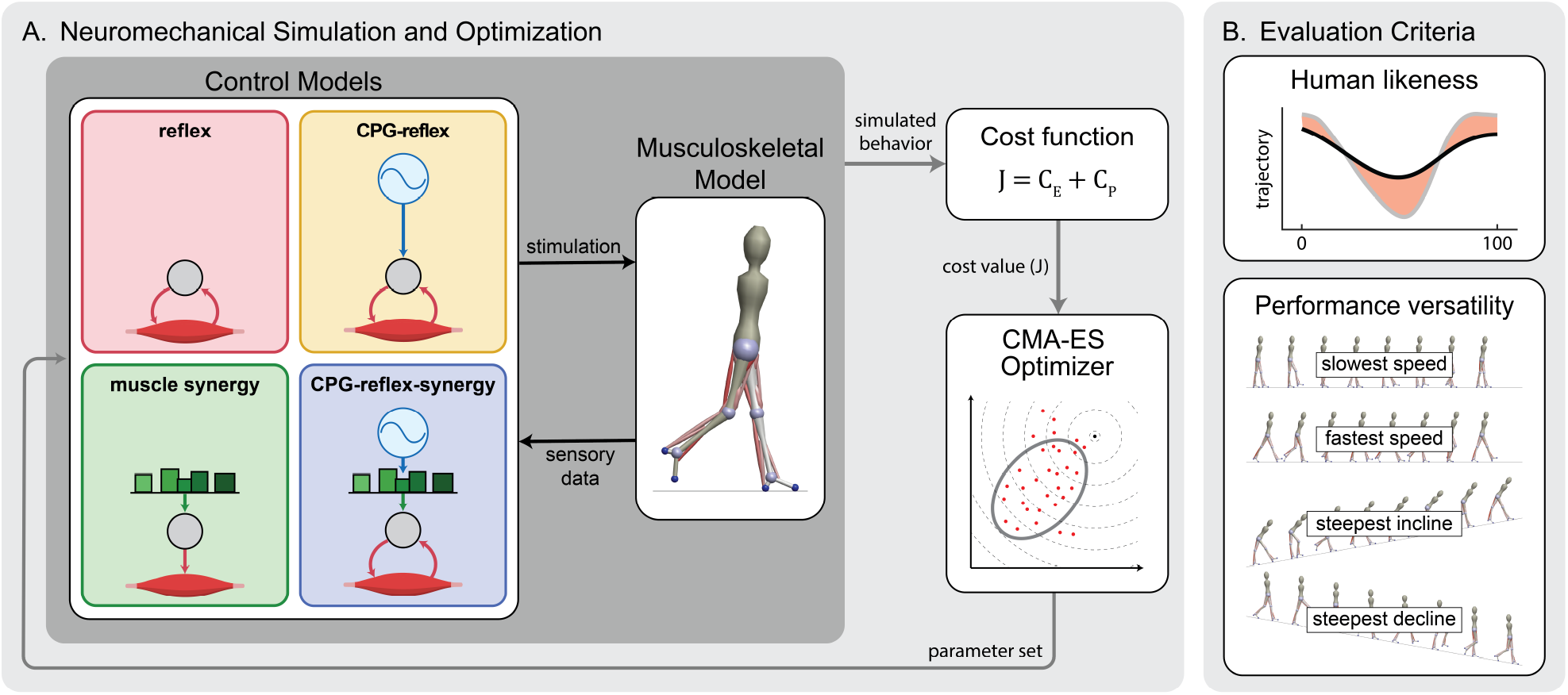
Simulation framework for comparative analysis. A) Neuromechanical simulation framework including neural control models, musculoskeletal model, and optimization. The control strategies include: reflex-based, CPG-reflex based, muscle synergy-based, and CPG-reflex-synergy based control strategies. B) Evaluation criteria include observing human-like biomechanical trends and performance versatility over speed–slope conditions.

### Existing neuromechanical locomotion control models

Neuromechanical simulations of locomotion have implemented a variety of control architectures reflecting different hypotheses about locomotor control. These models can be broadly categorized into reflex-based, CPG-reflex-based, muscle synergy-based, and hybrid architectures that integrate reflex, CPG, and synergy mechanisms. In this study, we selected one representative model from each category for implementation and comparison within a unified simulation framework (Table 1). The following subsections describe the selected models and the rationale for their selection.

**Table 1:** Representative neuromechanical locomotion control models implemented and compared in this study. (TD: transmission delays; ECC: excitation-contraction coupling)

| model | musculoskeletal model | compliant tendon | TD & ECC | optimization algorithm | objective | simulation platform | performance versatility |
| --- | --- | --- | --- | --- | --- | --- | --- |
| **Reflex-based**<br>(Song & Geyer, 2015a) | 3D, 7 segments,<br>22 muscles | ✓ | ✓ | CMA-ES | metabolic energy<br>(Umberger et al., 2003) | MATLAB/Simulink,<br>Simscape Multibody | $0.8\text{--}1.8\text{ m s}^{-1}$<br>$-13.5^\circ - +4.6^\circ$ |
| **CPG-reflex-based**<br>(Ogihara & Yamazaki, 2001) | 2D, 7 segments,<br>18 muscles | none | none | genetic algorithm | metabolic energy<br>(Hase & Yamazaki, 1997) | custom-built | $1.1\text{ m s}^{-1}$ |
| **Muscle synergy-based**<br>(Aoi et al., 2019) | 2D, 7 segments,<br>18 muscles | none | ✓ | manual-tuning | – | not reported | $1.2\text{--}1.6\text{ m s}^{-1}$ |
| **CPG-reflex-synergy-based**<br>(Di Russo et al., 2023) | 2D, 7 segments,<br>18 muscles | ✓ | ✓ | CMA-ES | metabolic<br>(Uchida et al., 2016) | SCONE | $0.3\text{--}1.9\text{ m s}^{-1}$ |

Reflex-based control models generate locomotion through physiologically inspired sensorimotor feedback pathways that transform proprioceptive signals into muscle activation without relying on an explicit CPG. Günther and Ruder (2003) proposed a walking control model rooted in the equilibrium point hypothesis (Feldman & Levin, 1995), in which muscle activation arises from deviations between actual and reference muscle lengths and velocities. A widely adopted model in this class is the sagittal-plane muscle-reflex model of Geyer and Herr (2010), which demonstrated human-like walking dynamics and muscle activation patterns with substantially improved agreement with experimental observations relative to earlier models. In this model, reflex pathways were designed to encode key leg functions for locomotion and constructed using delayed feedback from muscle force, length, and velocity signals (representing afferent inputs from Golgi tendon organs and muscle spindles), together with additional sensory signals such as trunk orientation and leg load. While the original study manually tuned the control parameters to produce human-like gait, subsequent studies showed that parameters optimized for metabolic energy expenditure can also produce realistic gait across a wide range of walking speeds (Song & Geyer, 2012). A large body of subsequent work adapted this model to generate different locomotor behaviors including walking and running (Song & Geyer, 2015b; Wang et al., 2012), extended it to broader speed ranges and terrains (Song & Geyer, 2012, 2015a, 2015b), generalized the framework to three-dimensional musculoskeletal models (Song & Geyer, 2015a; Wang et al., 2012), and applied it to simulations of elderly (Song & Geyer, 2018) or pathological gait (Ong et al., 2019). The Geyer and Herr model has also been integrated with additional control components, including higher-level control layers that modulate desired foot placement for robust walking (Song & Geyer, 2015a), controllers incorporating internal inverse models for voluntary swing-leg control (Ramadan et al., 2022), and feedforward components that improve robustness to step perturbations (Haeufle et al., 2018). The reflex-based controller has also been adopted as a baseline control architecture in the open-source neuromechanical simulation platform SCONE (Geijtenbeek, 2019), facilitating a wide range of model adaptations and applications (Fleischmann et al., 2025; Jones et al., 2024; Veerkamp et al., 2021; Waterval et al., 2023).

CPG-reflex-based control models generate locomotion through an internal rhythmic drive, typically implemented using neural oscillators, together with sensory feedback pathways that modulate the locomotor pattern. To our knowledge, the earliest forward simulations of human locomotion control using biologically inspired control models were introduced by Taga (1995), who demonstrated that stable walking patterns can emerge in a torque-driven multibody model through neural oscillators based on the formulation proposed by Matsuoka (1985). Building on this framework, Yamazaki, Hase, and colleagues developed what is, to our knowledge, the first neuromechanical model of human walking, integrating a motor control model with a musculoskeletal system in a forward dynamics simulation (Yamazaki et al., 1996). Their subsequent studies further investigated the integration and relative contributions of proprioceptive reflexes and CPGs in generating locomotion (Ogihara & Yamazaki, 2001), and later extended this line of work to what is likely the first three-dimensional neuromechanical simulation of human walking (Hase & Yamazaki, 2002). Later studies explored variations of this CPG-reflex architecture by modifying oscillator formulations or the interaction between rhythmic drive and sensory feedback. These include models employing alternative oscillator structures (Dzeladini et al., 2014; Paul et al., 2005; Severini & Muñoz, 2025), incorporating control modules resembling cerebellar error-correction mechanisms (Kim et al., 2011), or integrating CPG oscillators with the reflex pathways of Geyer and Herr (Dzeladini et al., 2014).

Muscle synergy-based control models generate muscle activation patterns through low-dimensional control signals that combine fixed synergy weights with time-varying activation pulses. Because muscle synergies primarily describe how a small number of activation modules are distributed across muscles rather than how these modules are generated, muscle synergy-based control architectures typically require an additional mechanism to produce the temporal activation signals.

In many implementations this rhythmic drive is generated by CPG-like mechanisms, while stability is maintained through simple feedback rules, often without explicitly modeling spinal reflex pathways. One of the earliest neuromechanical implementations of this concept was proposed by Jo and Massaquoi (2007), who constructed a walking controller in which the gait cycle was partitioned into discrete temporal windows associated with predefined muscle synergies. Aoi and colleagues developed a neuromechanical model in which temporal activation pulses with adjustable onset timing, duration, and amplitude drove predefined muscle synergies (Aoi et al., 2008). Subsequent studies within this framework incorporated additional control components, such as phase-resetting mechanisms (Aoi et al., 2010; Tamura et al., 2020) and speed regulation modules (Aoi et al., 2019), to examine how synergy-based control interacts with sensory feedback and task constraints in generating stable and adaptable locomotion.

CPG-reflex-synergy-based control models integrate rhythmic pattern generation, sensory feedback, and modular muscle coordination within a unified architecture. Such a hierarchical organization, in which central pattern generators provide rhythmic drive while sensory feedback and modular coordination shape muscle activation, is widely considered a plausible organization of spinal locomotor circuits in vertebrates (Danner et al., 2017). Recent neuromechanical simulations have explored implementations of this combined structure to integrate the complementary roles of CPG-based rhythm generation, reflex-mediated feedback control, and synergy-based muscle coordination within a single locomotion control framework (Di Russo et al., 2023).

## Methods

In this section, we first describe the four locomotion control models examined in this study, each representing one of the control strategies introduced above. We then present the unified neuromechanical simulation framework used to evaluate these controllers, which includes a 2-D sagittal-plane neuromusculoskeletal model and a standardized optimization procedure designed to assess gait performance in terms of human-likeness and versatility.

### Model selection and controller descriptions

We selected representative models from each of the four major control strategies: reflex-based, CPG-reflex-based, muscle synergy-based, and CPG-reflex-synergy-based approaches. Selection was guided by physiological plausibility, demonstrated capability to generate human-like and diverse locomotor behaviors, and how clearly each model represents the conceptual and historical foundation of its respective control architecture. For each model, we preserved the core control architecture while standardizing implementation details required for a consistent comparison within our simulation framework. Modifications to the control models were limited to those necessary to ensure compatibility with the shared musculoskeletal model and, when necessary, to maintain physiologically plausible gait behavior under these common conditions. The following subsections describe each selected model by briefly justifying its selection, summarizing the key elements of its control formulation, and specifying any implementation modifications made within our unified framework. Readers are referred to the original publications for detailed descriptions of each model (reflex-based (Song & Geyer, 2015a), CPG-reflex-based (Ogihara & Yamazaki, 2001), muscle synergy-based (Aoi et al., 2019), and CPG-reflex-synergy-based (Di Russo et al., 2023)), as well as to the open-source implementation provided with this study (*https://doi.org/10.5281/zenodo.20509551*).

#### Reflex-based model

The model of Song and Geyer (2015a) was selected to represent the reflex-based control strategy because it drives locomotion entirely by sensorimotor feedback, without an explicit CPG, designed in a manner that closely reflects the proprioceptive, vestibular, and cutaneous pathways of the nervous system. Beyond its physiological plausibility, this model has demonstrated a broad range of locomotor behaviors, including three-dimensional walking and turning, slope ascent and descent, stair ascent, obstacle avoidance (Song & Geyer, 2015a), and running (Song & Geyer, 2015b), and robustness against external perturbations (Song & Geyer, 2017), as well as extensions with direct clinical implications (Song & Geyer, 2018; Ton et al., 2024).

The controller is organized hierarchically into spinal and supraspinal layers. The spinal layer consists of ten modules divided between stance (M1–M5) and swing (M6–M10), each responsible for specific biomechanical functions such as compliant load support of the stance leg, effective swing toward target foot placement, and trunk balance. Muscle stimulation is generated through delayed feedback from muscle force, length, and velocity signals, representing afferent inputs from Golgi tendon organs and muscle spindles, along with additional sensory signals such as trunk orientation and leg load. These feedback pathways are parameterized through gains and offsets that shape muscle activation patterns. The supraspinal layer modulates foot placement based on hip position and velocity relative to the stance foot and coordinates gait phase transitions between stance and swing.

We implemented the 2-D version of this model (Song & Geyer, 2017, 2018; Ton et al., 2024) that omits hip adduction and abduction degrees of freedom while retaining all other control components, resulting in 64 optimization parameters.

#### CPG-reflex-based model

We selected the model of Ogihara and Yamazaki (2001) as representative of the CPG-reflex control strategy because it provides a clear and compact implementation of a rhythm generator coupled with sensory feedback within a neuromechanical simulation. This formulation directly extends the classical CPG-based locomotion framework introduced by Taga (1995), in which rhythmic neural activity interacts with body dynamics to produce stable gait. We also considered the model of Hase and Yamazaki (2002), but it relies on an optimization procedure at each simulation timestep to convert joint torques into muscle activations, which we considered physiologically implausible.

The controller follows a CPG-reflex architecture in which a rhythmic drive provides a baseline activation pattern, while sensory feedback modulates timing and amplitude at the motoneuron level. Rhythm generation is implemented using a Matsuoka oscillator, consisting of mutually inhibiting neurons with adaptation that produce alternating flexor–extensor activity (Matsuoka, 1985). Sensory feedback is incorporated through simplified neuron models representing muscle spindles and Golgi tendon organs, providing signals related to muscle length, velocity, and force.

Ground contact is represented through tactile signals based on foot–ground interaction, implemented as state-dependent inputs. For each muscle, an *α*-motoneuron integrates inputs from the oscillator, sensory pathways, contact signals, and a constant bias term to generate the final stimulation. Interactions between muscles spanning the same joint are also included, allowing excitatory and inhibitory coupling across muscles.

To improve robustness across walking conditions within the unified simulation framework, the oscillator parameters were included in optimization and a phase offset parameter *ϕ*_offset_ was introduced to define the initial states of the oscillators. As in the original formulation, oscillator dynamics were defined as:

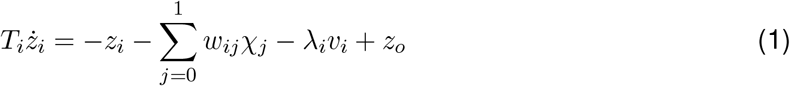

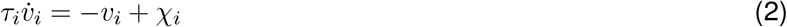

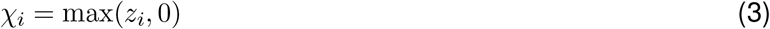

where *z*_*i*_ is the membrane potential of neuron *i* = 0, 1, *λ*_*i*_ and *v*_*i*_ represent the self-inhibition weight and state variable, *T*_*i*_ and *τ*_*i*_ are time constants, *w*_*ij*_ are synaptic connection weights, *z*_*o*_ is a constant input, and *χ*_*i*_ is the neuron output. For each simulation, the oscillator was first simulated offline using initial conditions *z*_0_(0) = *v*_0_(0) = *v*_1_(0) = 1 and *z*_1_(0) = 0. The initial states for the main simulation were then set as 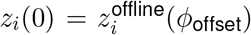 and 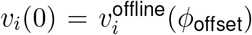. The oscillator parameters *T*_*i*_, *τ*_*i*_, *w*_*ij*_, *λ*_*i*_, *z*_*o*_, and *ϕ*_offset_ were included as tunable optimization variables. In total, the controller comprises 99 optimization parameters.

#### Muscle synergy-based model

The model of Aoi et al. (2019) was selected to represent the muscle synergy-based control strategy because it provides a direct implementation of low-dimensional temporal pulses that activate predefined muscle synergies. Alternative variants incorporating phase resetting at touchdown have been proposed (Aoi et al., 2010; Tamura et al., 2020); however, we selected this model as it includes speed control and demonstrated stable walking across speeds of approximately 1.2–1.6 m s^*−*1^, as well as running.

The controller is organized into a movement generator and a movement regulator. The movement generator produces muscle stimulation through five activation pulses distributed across the gait cycle, each associated with a predefined muscle synergy. Each pulse is characterized by its amplitude, onset timing, and duration, and contributes to muscle activation through weighted connections between pulses and individual muscles. The timing of these pulses is governed by a phase variable that progresses at a rate determined by gait period. The movement regulator complements this feedforward structure with feedback control, including a proportional–derivative controller for trunk balance acting on the hip flexors and glutei, and a proportional controller for speed regulation acting on the soleus and tibialis anterior. The total muscle stimulation is given by the combined output of the movement generator and regulator.

In the original formulation, the movement generator employs five square activation pulses defined by their onset timing and duration, producing piecewise-constant stimulation profiles across the gait cycle. In our implementation, these pulses resulted in less human-like kinematics and a reduced range of achievable walking speeds within the unified simulation framework. To address this, the square pulses (*p*_*i*_) were replaced with smoother bell-shaped activation curves using a polynomial pulse function:

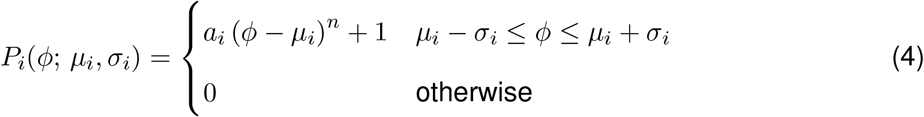

Where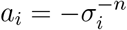 to normalize amplitude, *µ*_*i*_ = Φ_*i*_ + *σ*_*i*_ denotes the phase timing of peak amplitude where Φ_*i*_ is the onset phase of the pulse, and *σ*_*i*_ = Δ_*i*_*/*2 denotes the half-width of the pulse. The exponent *n* controls the curvature of the pulse. Based on preliminary simulations, *n* = 2 was adopted, as this parabolic profile produced more physiologically plausible joint kinematics and muscle activation patterns compared to the original square pulses. In total, the controller comprises 41 optimization parameters.

#### CPG-reflex-synergy-based model

We selected the model of Di Russo et al. (2023) to represent the CPG-reflex-synergy control strategy because it integrates a phase-resetting CPG with parameterized muscle synergies and spinal reflex pathways within a unified framework. This model extends prior CPG and reflex formulations by incorporating low-dimensional activation primitives while retaining physiologically motivated sensory feedback. The model generates a wide range of walking speeds without requiring external modulation.

The controller combines a phase-resetting CPG with muscle synergies and spinal reflex pathways to generate muscle stimulation. Rhythmic activity is produced by coupled oscillators, one for each leg, whose phases are reset at foot contact to maintain coordination with gait events. The oscillator phase drives a pattern formation layer that generates five bell-shaped activation primitives distributed across the gait cycle, each providing baseline stimulation to muscles through weighted connections. Sensory feedback is incorporated through spinal reflex pathways representing Ia, II, and Ib afferents as well as Renshaw cells, forming a network of sensory neurons, interneurons, and motoneurons that provide excitatory and inhibitory interactions across muscles. For each muscle, the final stimulation is determined by integrating inputs from the activation primitives and reflex pathways at the motoneuron level. In addition, a balance controller provides feedback-based modulation of hip muscles (HFL, GLU, and HAM) using a proportional–derivative control strategy.

The model was implemented following the original formulation without additional modifications. In total, the controller comprises 240 optimization parameters.

### Unified simulation framework and evaluation

We evaluated the ability of each control model to generate human-like gait and its performance across speeds and slopes using a unified neuromechanical simulation framework in the two-dimensional sagittal plane. In this framework, each controller drives a shared musculoskeletal model with nine muscles per leg, providing muscle stimulation inputs while interacting with identical biomechanical dynamics and simulation conditions. Gait behaviors were obtained through a standardized optimization procedure. The framework was implemented in MATLAB/Simulink (2024b) with the Simscape Multibody toolbox for rigid-body dynamics, and forward dynamics were computed using the ode15s solver.

#### Musculoskeletal model

We used the neuromechanical model of Song and Geyer (2018) with minor modifications. Specifically, to provide a more general and consistent framework across controllers, we adjusted the engagement threshold of the passive knee flexion torque preventing hyperextension to remove the earlier bias toward slight knee flexion during stance. In addition, we increased Achilles tendon compliance to improve tendon dynamics. The model represents an adult male with a height of 1.8 m and mass of 80 kg, and consists of a seven-segment rigid body system interacting with the ground via contact points at the heel and ball of each foot. The model is actuated by nine Hill-type muscle–tendon units per leg, each comprising a contractile element, a parallel elastic component, and a series elastic component. Physiological transmission delays and excitation–contraction coupling are included to represent neural and muscle dynamics. Detailed descriptions of segment properties, muscle–tendon parameters, activation dynamics, delay values, and the implemented model modifications are provided in Appendix A.

#### Optimization method

Control parameters were optimized using the Covariance Matrix Adaptation Evolution Strategy (CMA-ES) (Hansen, 2006). Each controller was tuned within a shared optimization framework, with a parameter space that included both controller-specific parameters and seven parameters defining the initial state of the musculoskeletal model (joint angles and horizontal velocity). The total number of control parameters differed across models: 64 for the reflex-based controller, 99 for the CPG-reflex-based controller, 41 for the muscle synergy-based controller, and 240 for the CPG-reflex-synergy-based controller.

Optimization was performed using a multi-stage objective designed to ensure stable and physiologically plausible gait. The cost function was defined as:

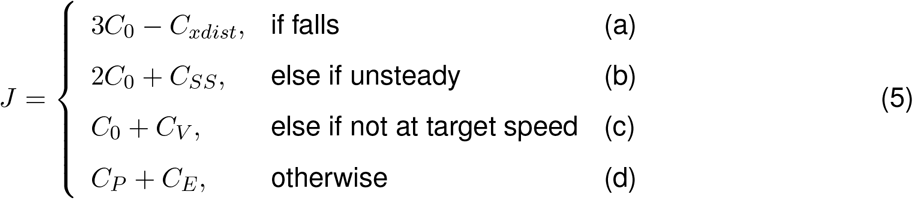

where *C*_0_ = 10^4^ ensures prioritization of earlier stages. The first stage promotes forward progression by maximizing traveled distance (*C*_*xdist*_). The second stage enforces steady walking, where (*C*_*SS*_) is defined as the summed differences in joint positions at successive heel strikes; steady gait is achieved when six consecutive steady-state steps are observed. The third stage enforces target speed tracking, defined as *C*_*v*_ = *max* (|*v*_avg_ |*− v*_tgt_ *−* 0.02, 0). The final stage minimizes joint limit violations (*C*_*P*_; hip: −25° to +135°, knee: −3° to 55°, ankle: −65° to 20°) and muscle effort 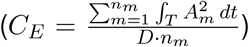, where muscle effort is computed as the distance-normalized integral of squared muscle activations across all muscles over the steady-state steps. Thus, a successfully optimized simulation trial begins from an initial pose and forward velocity, with the controller generating muscle stimulations based on its internal states and sensory feedback to drive motion, and ultimately converges toward a steady walking pattern with small step-to-step deviations, average speed within the target range, and minimized joint-limit violations, and distance-normalized muscle activations.

Optimization was initialized with an identity covariance matrix and terminated when improvements in the cost function fell below 3% over the last 200 iterations. A standard optimization consisted of 800 iterations. All parameters were normalized to an initial value of 1, and the initial sampling variance (*σ*) was set to 0.1 when exploring new conditions and reduced to 0.01 for continued optimization within the same condition. Computations were performed on a local multi-core workstation and Northeastern University’s high-performance computing cluster (Explorer), with typical runtimes of 3–10 hours locally and 6–15 hours on the cluster, depending on the controller.

#### Simulation protocol

Nominal walking solutions were obtained by optimizing controller parameters to produce gait with minimum muscle effort at a target speed of *v*_tgt_ = 1.2 m s^*−*1^ on level ground. To initialize this process, parameters were first optimized to minimize kinematic deviation between simulated and experimental gait data (Rose & Gamble, 2006), using a modified multi-stage cost function in which the final-stage objective (Eq. 5(d)) was replaced with the root mean square error of joint kinematics (see Appendix B). The resulting parameter set was then used as an initialization for further optimization under nominal conditions using the original multi-stage objective, producing stable, minimum-effort gait without reference data.

Performance across speeds and slopes was evaluated by systematically extending optimization from the nominal solution. Target walking speed (*v*_tgt_) was incrementally increased and decreased in steps of 0.2 m s^*−*1^ until stable gait could no longer be achieved. For each successfully optimized speed condition, the resulting parameter set was used to initialize optimization across slope variations. Incline and decline slopes were increased in increments of 1° until stable gait was no longer attainable. A speed–slope condition was considered achievable if the optimization reached the final stage of the cost function (i.e., minimization of *C*_*E*_ + *C*_*P*_), at which point optimization was terminated to reduce computational cost. This procedure was repeated multiple times to ensure that observed performance limits reflected the capabilities of each control model rather than convergence to local minima.

To further characterize performance trends, full optimization of the final-stage objective was performed for level ground across the range of achievable speeds and for slope conditions ranging from *−*10° to 10° at *v*_tgt_ = 1.2 m s^*−*1^. Exploration of speed–slope conditions continued until smooth unimodal performance trends were obtained for each control model.

#### Evaluation

We evaluated each control model along two dimensions: human-like gait characteristics and locomotor versatility. Human-like gait characteristics were assessed by comparing simulated kinematics, kinetics, and muscle activations against experimental reference data under nominal walking conditions, as well as by examining biomechanical (e.g., stride length) and energetic trends across walking speeds and slope conditions. Locomotor versatility was evaluated as the range of speed–slope conditions over which each controller could generate stable gait under the standardized optimization framework. Conditions exhibiting a flight phase (i.e., running gaits) on level ground were excluded from this analysis. Together, these evaluations quantify both the fidelity of simulated gait to human data and the adaptability of each control architecture across task conditions.

## Results

We first present results on human-like gait characteristics, including nominal gait patterns and trends across speeds and slopes, and then examine locomotor versatility across speed–slope conditions. Simulation animations for nominal gaits and representative speed–slope conditions are provided in the Supplementary Video (*https://youtu.be/suMJIfZe5k?si=GeTNjrnSL4SfAJ-T*).

### Human-like gait characteristics

#### Nominal gait patterns

The reflex-based and CPG-reflex-synergy models most closely reproduced human gait characteristics under nominal walking conditions. Fig. 2 shows the kinematics, kinetics, and muscle activations across the gait cycle for each controller when optimized using Eq. 5 at 1.2 m s^*−*1^. All models successfully generated stable walking. However, all models exhibited straightened or hyperextended knees during midstance, accompanied by mismatches in knee joint torque during stance. The reflex-based and CPG-reflex-synergy models produced more human-like gait kinematics, which is also evident in the simulation animations (Supplementary Video). The bar plots quantify deviations from experimental data using cross-correlation (1 *− R*) and root mean square error (RMSE) across joints, ground reaction forces, and muscle activations, where lower values indicate better agreement. Deviations were largest in muscle activations, particularly due to excessive hamstring activity and reduced rectus femoris activation. Overall, the reflex-based and CPG-reflex-synergy models showed the best agreement across kinematics, kinetics, and muscle activations in both RMSE and cross-correlation (1 *− R*), except for reduced correlation (higher 1 *− R*) in muscle activations for the CPG-reflex-synergy model.

**Figure 2:**
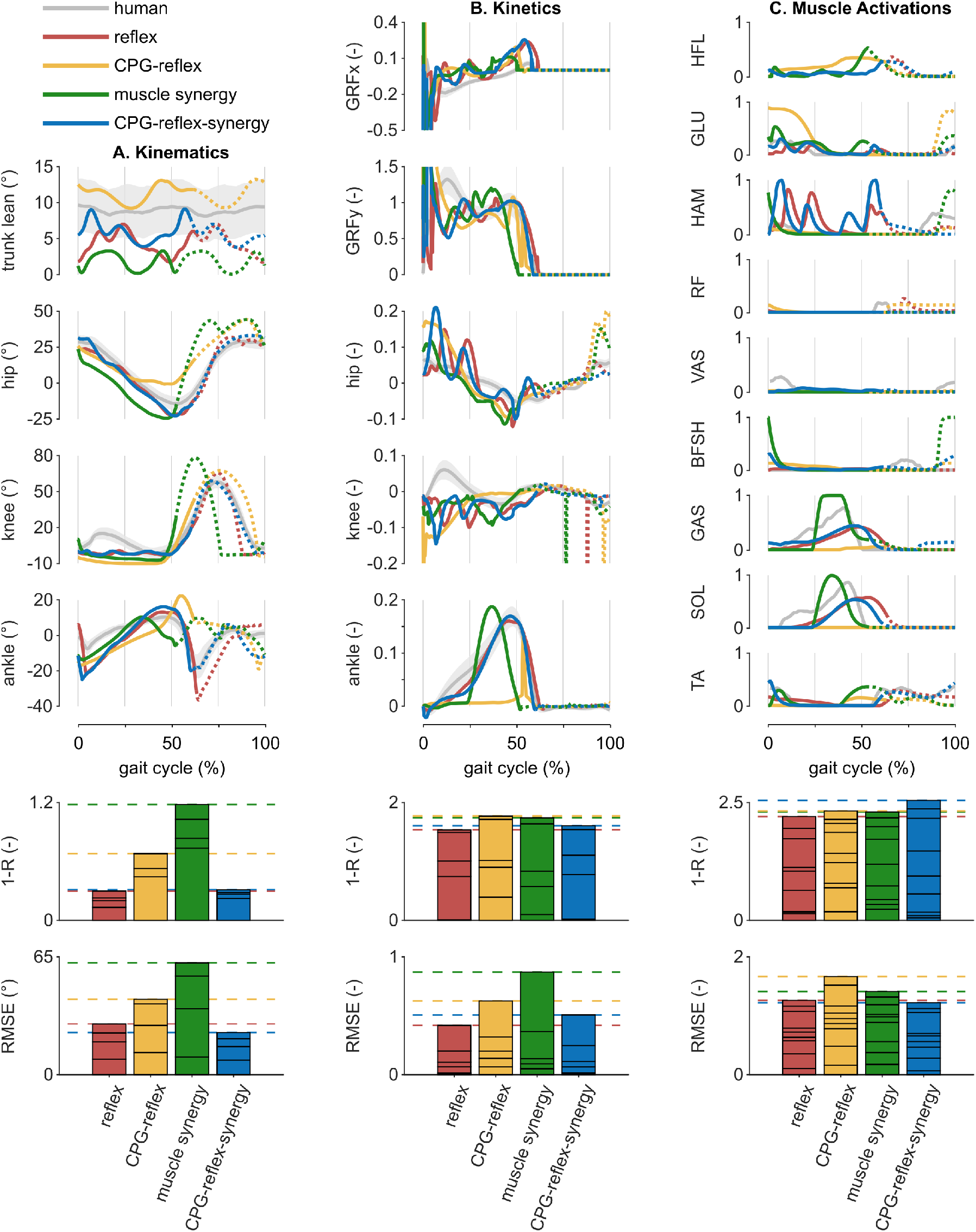
Kinematics, kinetics, muscle activations, and deviation of simulated to reference data for walking at 1.2 m s^*−*1^. A) human kinematics (Rose & Gamble, 2006), B) kinetics (Camargo et al., 2021), and C) muscle activations (Perry & Burnfield, 2010) are shown in gray. Ground reaction forces are normalized by body weight, and joint torques are normalized by body weight times leg length. Solid lines display trajectories during stance, while dashed lines represent swing. Bar plots show deviation of simulated gait data from reference data. Each stacked bar shows cross-correlation (1 *−R*) and root mean square error per joint, ground reaction force, or muscle activation with horizontal dashed lines noting the summed metric. The cross-correlation coefficient (*R*) is computed allowing up to a 20% time shift, and is presented as 1 *−R* so that larger values indicate greater deviation.

#### Trends across speeds and slopes

We compare stride length and metabolic energy trends across walking speeds on level ground and across slope conditions at a fixed speed of 1.2 m s^*−*1^, and report the corresponding optimal walking speed and slope for each energy and fatigue cost of transport metric. Muscle effort is reported as fatigue of transport (FOT), based on squared muscle activation as a proxy for fatigue (Song & Geyer, 2018), and is compared alongside two metabolic cost of transport metrics based on different muscle metabolic models (COT_U_: Umberger (2010); COT_B_: Bhargava et al. (2004)). Optimal values are obtained by fitting fourth-order polynomials to each trend and identifying their minima. Step length and metabolic cost of transport from experimental data are included for comparison.

The reflex-based and CPG-reflex-synergy models best captured human trends in stride length and energy across walking speeds on level ground. All models except the CPG-reflex model reproduced the characteristic increase and plateau in stride length with speed (Fig. 3). The reflex-based model was the most energy efficient, followed by the CPG-reflex-synergy model, and both showed consistent trends across energy metrics. In contrast, the CPG-reflex and muscle synergy models exhibited inconsistent energy trends, aligning with experimental data in COT_U_ but not in FOT or COT_B_. Consistent with these trends, the reflex-based model produced the closest optimal walking speed to experimental values (Table 2).

**Figure 3:**
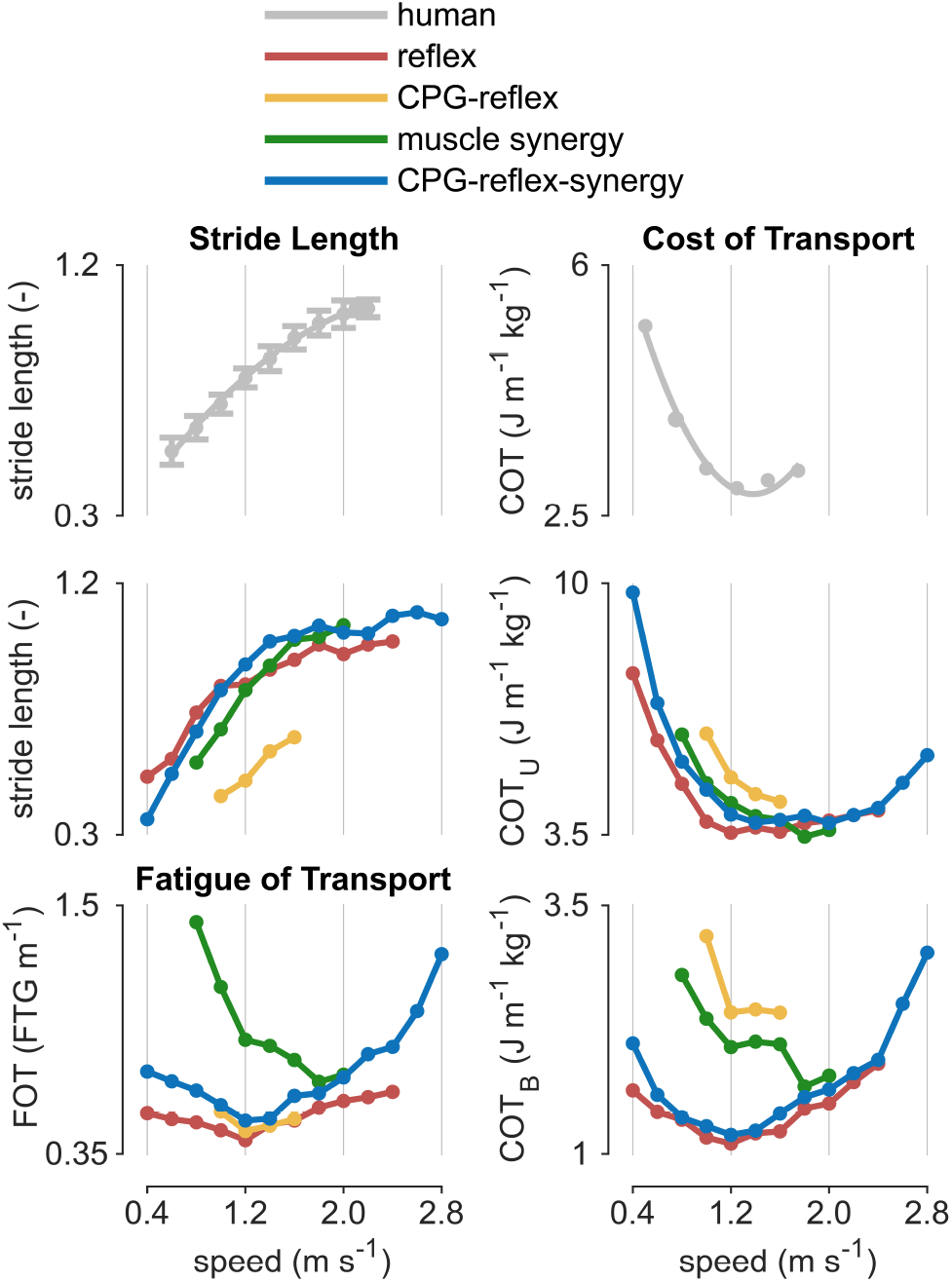
Biomechanical and energy trends of walking at different speeds on level ground. Stride length is normalized by subject height. Simulated results include fatigue of transport (FOT) and metabolic cost of transport (COT). Reference human data (top row) for stride length and COT are from Hirasaki et al. (1999) and Browning et al. (2006), respectively.

**Table 2:** Optimal walking speed on level ground conditions.

| control strategy | FOT | COT<sub>U</sub> | COT<sub>B</sub> |
| --- | --- | --- | --- |
| human data | 1.2-1.4 m s<sup>-1</sup> (Fritz & Lusardi, 2009) |  |  |
| reflex | 1.15 m s<sup>-1</sup> | 1.41 m s<sup>-1</sup> | 1.23 m s<sup>-1</sup> |
| CPG-reflex | 1.24 m s<sup>-1</sup> | > 1.60 m s<sup>-1</sup> | 1.27 m s<sup>-1</sup> |
| muscle synergy | 1.90 m s<sup>-1</sup> | 1.88 m s<sup>-1</sup> | 1.92 m s<sup>-1</sup> |
| CPG-reflex-synergy | 1.18 m s<sup>-1</sup> | 1.49 m s<sup>-1</sup> | 1.05 m s<sup>-1</sup> |

Across slope conditions, the reflex-based and CPG-reflex-synergy models most consistently reproduced experimentally observed stride length and energy trends (Fig. 4). It is worth noting that trends across slopes were more irregular than those on level ground, reflecting the increased difficulty of optimization and the presence of local minima. Both models captured the characteristic peak in stride length at moderate incline (approximately 3°), whereas the CPG-reflex and muscle synergy models did not exhibit human-like stride length patterns across slopes. Although the muscle synergy model produced optimal slope values closest to experimental data for some energy metrics (Table 3), it remained the least energy efficient across all slope conditions. In contrast, the reflex-based and CPG-reflex-synergy models achieved lower energy cost across slopes while producing optimal slope conditions consistent with experimental observations.

**Figure 4:**
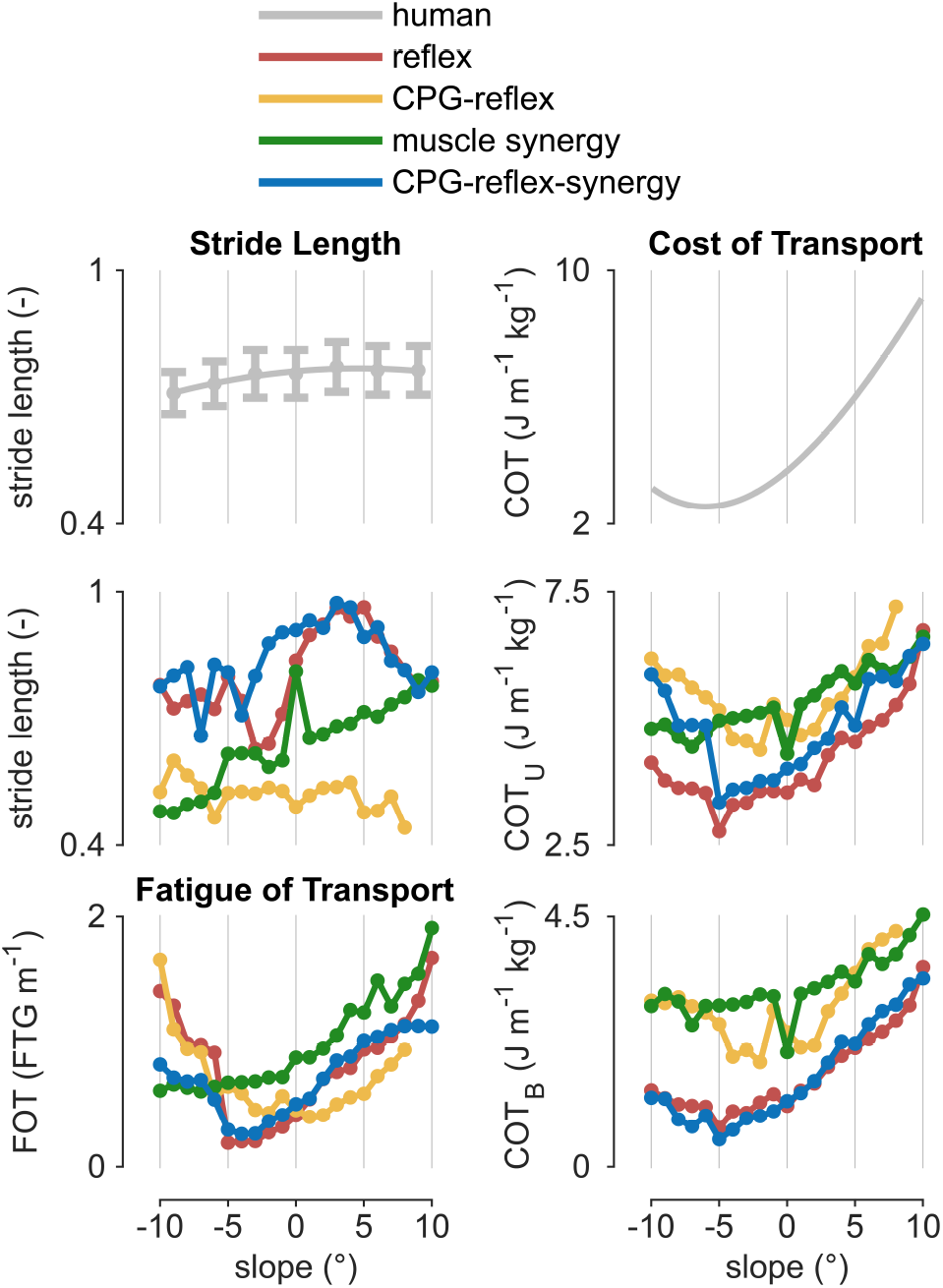
Biomechanical and energy trends of walking across different slope conditions at 1.2 m s^*−*1^. Stride length is normalized by subject height. Simulated results include fatigue of transport (FOT) and metabolic cost of transport (COT). Reference human data (top row) for stride length and COT are from Franz et al. (2012) and Lankford et al. (2020), respectively.

**Table 3:** Optimal slope condition for walking at 1.2 m s^*−*1^.

| control strategy | FOT | COT<sub>U</sub> | COT<sub>B</sub> |
| --- | --- | --- | --- |
| human data | -6.00° (Lankford et al., 2020) |  |  |
| reflex | -2.12° | -4.95° | -5.35° |
| CPG-reflex | -0.18° | -1.09° | -1.45° |
| muscle synergy | -7.37° | -7.37° | -4.14° |
| CPG-reflex-synergy | -3.13° | -2.32° | -4.95° |

### Versatility across speeds and slopes

The CPG-reflex-synergy model achieved the greatest versatility across walking speeds and slopes, followed by the reflex-based model, whereas the muscle synergy and CPG-reflex models showed substantially more limited coverage (Fig. 5). Both the CPG-reflex-synergy and reflex-based models produced very wide speed ranges (0.4–2.8 m s^*−*1^ and 0.4–2.4 m s^*−*1^, respectively), and were able to achieve even faster locomotion with running gaits, although these conditions were excluded from this analysis. In contrast, the muscle synergy and CPG-reflex models were restricted to narrower speed ranges. Similarly, for slopes, the CPG-reflex-synergy model achieved the broadest range of inclines and declines, while the reflex-based model showed comparable but slightly reduced coverage, whereas the muscle synergy and CPG-reflex models again exhibited more limited adaptability.

**Figure 5:**
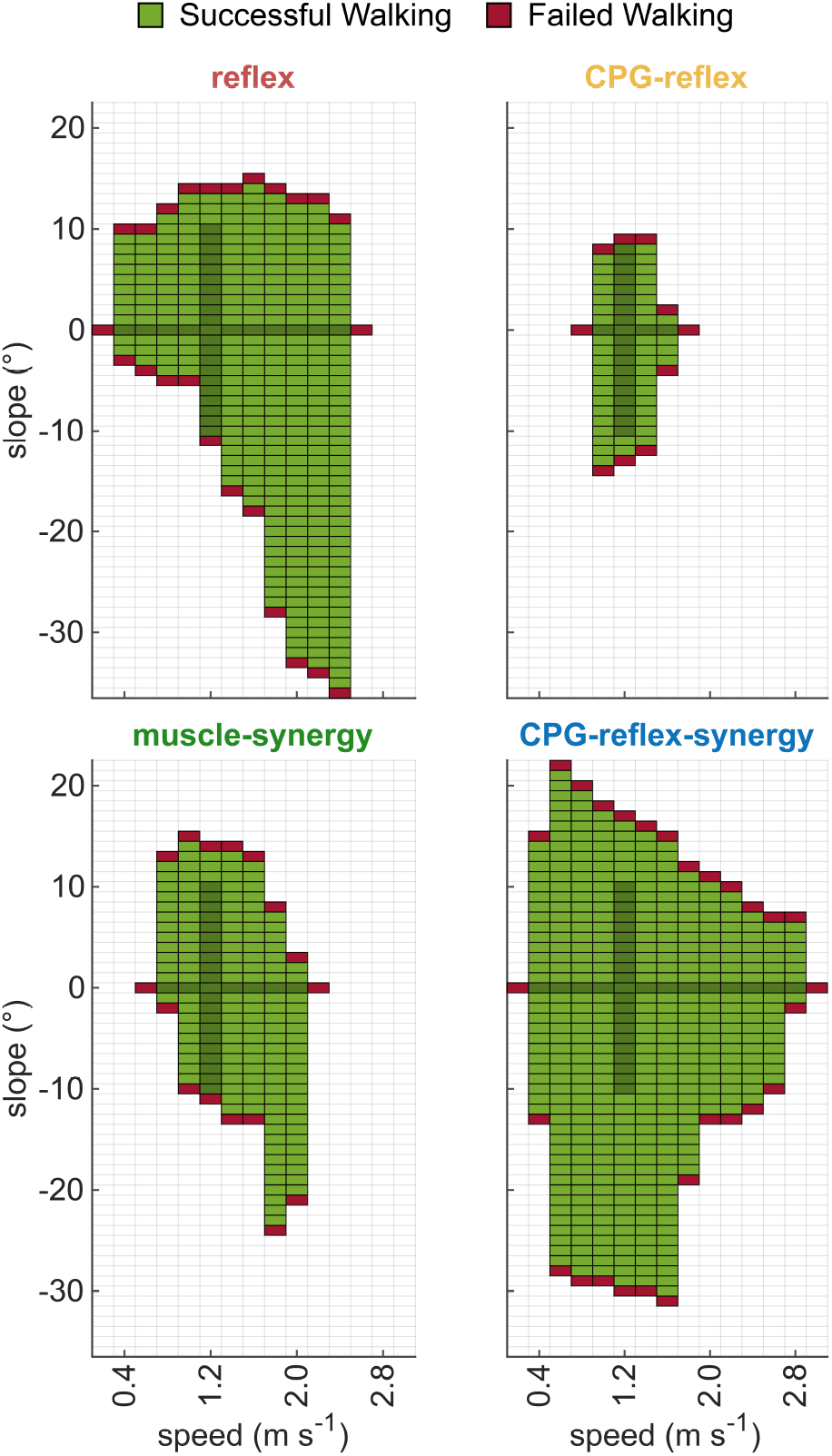
Performance versatility of each controller across walking speeds and slopes. Each grid cell represents a simulation at a given speed–slope condition. Successful walking is indicated in green, while red denotes unsuccessful cases. Darker green cells indicate conditions that were fully optimized and included in Figs. 3 and 4.

### Comparison with previously reported model behaviors

The implemented controllers reproduced the characteristic kinematic, kinetic, and muscle activation patterns reported in their original publications and achieved a wider range of locomotion behaviors, while enabling evaluation under a substantially broader and unified set of locomotion conditions (Table 4). Readers are referred to the original studies for more detailed locomotion behaviors.

**Table 4:** Comparison of implemented controllers with previously reported model behaviors.

| control strategy | previously reported behavior | behavior in this study | previously reported range | range in this study |
| --- | --- | --- | --- | --- |
| reflex | Kinematics matched human gait well, except for high peak hip flexion, early knee extension and a more plantarflexed ankle during stance. Low knee torque, early peak ankle torque, and high hip with low ankle muscle activity observed. (Song & Geyer, 2015a). | Similar overall gait pattern with more natural hip and ankle motion during late stance, but with straightened knees through midstance and increased ankle plantarflexion during early stance. Increased HAM activity remained, but more natural ankle torque observed. | $0.8\text{--}1.8\text{ m s}^{-1}$;<br>$-13.5^\circ$ to $4.5^\circ$<br>slopes | $0.4\text{--}2.4\text{ m s}^{-1}$;<br>$-35^\circ$ to $14^\circ$<br>slopes |
| CPG-reflex | Limited ankle motion, straightened knee during stance throughout the gait cycle. High hip torque correlation to human data, but low peak knee and ankle torque produced. Muscle activation not reported. Muscle tension reported as comparable; refer to the original paper (Ogihara & Yamazaki, 2001). | Similar knee and hip patterns with straightened knees through midstance; ankle kinematics differed but still showed limited range of motion, low knee and ankle torque, and minimal GAS activation. | $1.1\text{ m s}^{-1}$ on level ground | $1.0\text{--}1.6\text{ m s}^{-1}$;<br>$-13^\circ$ to $8^\circ$<br>slopes |
| muscle synergy | Straightened knee during stance, high ankle plantarflexion, and early hip flexion during swing. Weaker joint torques observed for all three joints. Increased BFSH, VAS, GAS, and TA activations reported (Aoi et al., 2019). | Similar kinematic features, including straightened knees through midstance, high ankle plantarflexion, and early hip flexion during swing. Peak joint torques matched human data, but temporally did not align. Higher activations for BFSH, GAS, and TA remained. | $1.2\text{--}1.6\text{ m s}^{-1}$ walking;<br>$1.6\text{--}2.0\text{ m s}^{-1}$ running | $0.8\text{--}2.0\text{ m s}^{-1}$ walking; $-23^\circ$ to $14^\circ$ slopes |
| CPG-reflex-synergy | Kinematics and vertical GRF matched human gait well, except with high peak hip flexion during swing, excessive knee flexion during mid-to-late stance, and more dorsiflexed ankle posture during early stance and early swing. Joint torques and horizontal ground reaction force not reported. Low GAS and SOL reported. (Di Russo et al., 2023). | High peak hip flexion during swing was not observed; straightened knees through midstance and increased ankle plantarflexion during early stance were observed. Increased activity for hip muscles. | $0.3\text{--}1.9\text{ m s}^{-1}$ on level ground | $0.4\text{--}2.8\text{ m s}^{-1}$;<br>$-30^\circ$ to $21^\circ$<br>slopes |

## Discussion

This study shows that specific implementations of locomotion control architectures can meaningfully shape both the human-likeness and versatility of neuromechanical simulations under controlled biomechanical and computational conditions. By implementing four representative controllers within a unified musculoskeletal model, optimization procedure, and evaluation framework, we isolated differences attributable to control organization rather than to differences in body mechanics or simulation protocol. The CPG-reflex-synergy model of Di Russo et al. (2023) achieved the strongest overall performance across the evaluated criteria, producing human-like gait patterns and the widest range of walking speeds and slopes. The reflex-based model of Song and Geyer (2015a) showed similarly strong human-likeness and competitive versatility, whereas the muscle synergy-based model of Aoi et al. (2019) and the CPG-reflex-based model of Ogihara and Yamazaki (2001) showed less realistic gait patterns and more restricted adaptability. These results indicate that the organization of neural control, as instantiated in specific model formulations, is a major determinant of simulated locomotor behavior.

The two best-performing models shared a common feature: their control signals were grounded in the mechanical state of the body and its interaction with the environment. In the reflex-based model, muscle activation is continuously modulated by sensory feedback related to muscle state, limb loading, and trunk orientation, while stance and swing reflex circuits are coordinated through gait events such as ground contact. In the CPG-reflex-synergy model, muscle activation is continuously modulated through reflex pathways, in addition to the CPG-generated rhythm being synchronized with body–environment interaction through phase resetting at heel-strike. Both models also incorporate interlimb coordination, through contralateral leg feedback in the reflex-based model and bilateral CPG coupling in the CPG-reflex-synergy model. In contrast, the CPG-reflex and muscle synergy models rely more heavily on internally generated timing, with less explicit sensory anchoring of the locomotor pattern and less direct coupling between limbs. These differences may explain why the reflex-based and CPG-reflex-synergy models produced stable gait across broader ranges of speeds and slopes: successful walking requires not only generating rhythmic activation patterns, but also continuously modulating those patterns according to body state and mechanical events. This control requirement is captured in forward-dynamic simulations, where gait must emerge from transient initial conditions and remain stable despite step-to-step variability. In contrast, trajectory optimization approaches are well suited for identifying optimal periodic motions but do not typically test how a neural controller stabilizes gait from transient or imperfect states unless feedback, perturbations, or noise are explicitly modeled (De Groote & Falisse, 2021).

Several additional observations emerged from the unified comparison. Notably, the two controllers that produced gait less consistent with experimental observations were originally developed without some key neuromechanical features, such as compliant tendons and physiological transmission delays (Table 1), highlighting the importance of neuromechanical modeling detail in evaluating locomotion control architectures. In addition, controller dimensionality varied substantially across models, with 41, 64, 99, and 240 parameters for the muscle synergy-based, reflex-based, CPG-reflex-based, and CPG-reflex-synergy-based controllers, respectively. While the greater dimensionality of the CPG-reflex-synergy-based controller may provide increased representational capacity, the reflex-based controller nevertheless achieved comparable overall performance despite using substantially fewer parameters.

The results of this study should be interpreted as comparisons of specific model implementations, not as definitive judgments on the underlying biological control hypotheses. Reflexes, CPGs, and muscle synergies each represent broad classes of mechanisms that can be instantiated in many different ways. The weaker performance of the CPG-reflex and muscle synergy models in this study therefore does not imply that CPGs or synergies are unimportant for human locomotion. Rather, it indicates that the particular formulations tested here were less effective under the shared framework and evaluation criteria. Different implementations, such as CPG-reflex models with richer sensory feedback, synergy models with stronger event-based timing regulation, or hybrid controllers with alternative spinal circuit representations, could produce different results. The present study should therefore be viewed as a controlled comparison of representative architectures and a tool for identifying useful design principles, rather than as a final ranking of biological control mechanisms.

More broadly, the limited versatility observed in some controllers should also be interpreted in the context of the level of control these models were designed to represent. Most biologically inspired locomotion models, including those examined here, primarily focus on spinal-level mechanisms responsible for generating and regulating steady-state locomotion through reflex pathways, rhythmic circuits, and modular muscle coordination. From this perspective, reduced adaptability across broader speed and slope conditions does not necessarily invalidate the underlying control hypothesis, as flexible locomotor behaviors in humans likely depend on additional supraspinal processes involved in regulating task goals, balance, gait transitions, and environmental adaptation. In the present framework, such higher-level adaptation was approximated through re-optimization of controller parameters across conditions rather than through explicit hierarchical modulation during locomotion itself. The present results therefore motivate future neuromechanical studies that model locomotion beyond nominal steady gait to investigate more holistic control architectures that couple spinal locomotor mechanisms with supraspinal systems capable of context-dependent modulation and task-level regulation.

### Limitations

Our unified simulation framework has several limitations related to the simplified musculoskeletal model and optimization framework. The musculoskeletal model includes several simplifications that would affect the resulting behaviors of different control models: it is restricted to the sagittal plane, includes a reduced set of muscles, represents the upper body as a single rigid segment, and does not include a toe segment. Nevertheless, the selected framework is appropriate for the present comparison because many of the original neuromechanical control models were either proposed using similar musculoskeletal formulations or later adapted to comparable reduced-order models by the original authors. The optimization framework also imposes assumptions regarding locomotor goals and optimality principles. In particular, muscle effort (computed as the summed integral of squared muscle activations) was used as the primary final-stage objective. Although this objective is widely used in musculoskeletal simulations and has been argued to produce human-like gait patterns (Ackermann & Van den Bogert, 2010; Song & Geyer, 2018), alternative objectives such as metabolic energy expenditure (Koelewijn et al., 2019; Song & Geyer, 2018) or multi-objective formulations (Veerkamp et al., 2021) may produce different locomotor behaviors and alter the relative performance of the controllers. However, when optimized to better match experimental gait kinematics (Appendix B), the CPG-reflex-based and muscle synergy-based models still showed limited improvement in reproducing the reference kinematics, suggesting limitations of those control formulations in generating human-like gait within the present framework beyond the specific objective formulation used in the main analysis. Another limitation is that, despite repeated optimization attempts, the high-dimensional non-convex optimization landscape likely contains local minima, particularly for slope conditions where irregular trends remained apparent (Fig. 4), and the obtained solutions may therefore not represent globally optimal locomotor behaviors.

Another potential concern is that the modeling and framework choices used in the present study may bias the comparison toward certain control models, particularly the reflex-based model, because the musculoskeletal model and simulation environment were adapted from prior reflex-based modeling studies. We sought to reduce this concern by applying the same musculoskeletal dynamics, optimization procedure, simulation protocol, and evaluation metrics to all controllers, and by limiting implementation changes to those required for compatibility or physiologically plausible performance. We also modified several aspects of the original framework to reduce behavior-specific tuning and incorporate more physiologically realistic musculoskeletal dynamics, including removing a passive knee torque threshold that previously prevented full knee extension and incorporating more compliant Achilles tendon properties and more detailed activation dynamics. These changes likely contributed to several gait differences from prior reflex-based studies, such as the excess knee extension observed during early stance. In addition, kinematic-fitting optimization was used to obtain suitable initial parameter sets for the final optimization stage across all controllers, and the full optimization procedure was repeated multiple times to reduce sensitivity to local minima within practical computational constraints. Moreover, the implemented controllers achieved broader speed and slope ranges than those reported in their original studies, suggesting that the unified framework explored the capabilities of all control models to a substantial extent. Nevertheless, no unified comparison can be completely independent of modeling assumptions, and the present results should therefore be interpreted in the context of the selected representative models, shared body mechanics, and chosen evaluation criteria. To further support transparency and reproducibility, we publicly share the unified simulation framework and controller implementations, allowing future studies to reproduce, extend, and critically evaluate the present comparison under alternative modeling assumptions and optimization strategies.

### Future Directions

The public framework developed in this study provides a foundation for systematic and extensible comparison of neuromechanical locomotion control models. Future work can implement additional variants within each control class to determine which representations of reflex pathways, CPG dynamics, synergy structures, sensory feedback, and phase regulation are most important for generating human-like gait. The framework can also be extended to investigate other performance dimensions and applications, including robustness to external perturbations, gait transitions, pathological gait features, motor adaptation and learning, rehabilitation interventions, and interactions with assistive devices. Beyond controller comparisons, the framework may also be used to investigate how musculoskeletal morphology, muscle parameters, tendon compliance, objective functions, optimization algorithms, and physics solvers influence simulated locomotor behavior. Such studies would help distinguish conclusions that are specific to particular controller implementations from those that reflect broader interactions among neural control, body mechanics, and task constraints.

More broadly, to investigate locomotor behaviors beyond nominal steady gait, future neuromechanical studies may benefit from more holistic models that explicitly integrate spinal control primitives with supraspinal modulation. Rather than treating changes in speed, slope, or task condition as separate optimization problems, such hierarchical models could include explicit mechanisms for task-level control, sensory state estimation, balance regulation, motor adaptation, and context-dependent modulation of spinal circuits. Such models would provide a stronger computational bridge between spinal locomotor mechanisms, supraspinal regulation, and whole-body biomechanics. Because designing such higher-level control mechanisms manually may become increasingly difficult as locomotor tasks and environments grow more complex, reinforcement learning and related data-driven approaches may provide useful tools for learning supraspinal modulation policies over physiologically grounded spinal control architectures (Song et al., 2021). By enabling systematic comparison of alternative control architectures under shared conditions, the present framework provides a foundation for such investigations and may help guide the development of more physiologically grounded models of human locomotor control.

## Appendix A Musculoskeletal Parameters

The musculoskeletal model is visualized in Fig. A1. The skeletal model properties (Table A1) are derived from physiological measurements (Winter, 2009). The muscle-tendon parameters (Table A2) and musculoskeletal attachment parameters (Table A3) are set on physiological data (Miller, 2014; Song & Geyer, 2015a, 2018). Neural delays are included in the simulation representing transmission from the spinal cord to the hip, knee, and ankle muscles with values of 2.5 ms, 5 ms, and 10 ms, respectively. A 30 ms delay is applied for signals from the supraspinal system to spinal cord. From our previous standard simulation framework (Song & Geyer, 2015a, 2018), the Achilles tendon properties have been updated to be more compliant, better matching reported tendon kinematics (Arnold et al., 2013) and the engagement threshold of the passive knee flexion torque is also relaxed from 175° to 180°. Readers are referred to the code shared for full details on the musculoskeletal implementation.

**Figure A1:**
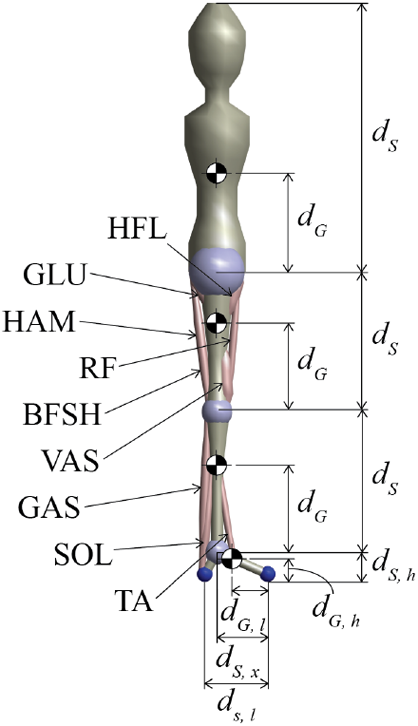
Musculoskeletal model used in simulation framework. The nine lower limb muscles are labeled on the left, while limb lengths (*d*_*S*_) and center of mass positions (*d*_*G*_) are depicted on the right.

**Table A1:** Skeletal segment parameters.

|  | HAT | Thigh | Shank | Foot |  |  |
| --- | --- | --- | --- | --- | --- | --- |
|  |  |  |  | l | h | x |
| $d_S (cm)$ | 84.6 | 44.10 | 44.28 | 20 | 7.02 | 16 |
| $d_G (cm)$ | 35 | 28.24 | 28.24 | 14 | 6.14 | - |
| $m_S (kg)$ | 54.24 | 8 | 3.72 | 1.16 | | |
| $\Theta_z (kg \cdot m^2)$ | 2.5 | 0.15 | 0.05 | 0.005 | | |

**Table A2:** Muscle-tendon parameters.

|  | HFL | GLU | HAM | RF | VAS | BFSH | GAS | SOL | TA |
| --- | --- | --- | --- | --- | --- | --- | --- | --- | --- |
| $F_{max} (kN)$ | 2 | 4 | 2 | 0.85 | 5 | 0.35 | 2 | 4 | 0.8 |
| $v_{max} (l_{opt} s^{-1})$ | 12 | 12 | 12 | 12 | 12 | 12 | 12 | 6 | 12 |
| $l_{opt} (cm)$ | 11 | 11 | 10 | 8 | 8 | 12 | 5 | 4 | 6 |
| $l_{slack} (cm)$ | 10 | 13 | 31 | 35 | 23 | 10 | 40 | 26 | 24 |
| $\varepsilon_{ref} (-)$ | 0.04 | 0.04 | 0.04 | 0.04 | 0.04 | 0.04 | 0.10 | 0.10 | 0.04 |

**Table A3:** Musculoskeletal attachment parameters.

|  | Hip |  |  |  | Knee |  |  |  |  | Ankle |  |  |
| --- | --- | --- | --- | --- | --- | --- | --- | --- | --- | --- | --- | --- |
|  | HFL | GLU | HAM | RF | HAM | RF | VAS | BFSH | GAS | GAS | SOL | TA |
| $r_0 (cm)$ | 8 | 8 | 8 | 8 | 5 | - | - | 4 | - | - | - | - |
| $r_{max} (cm)$ | - | - | - | - | - | 6 | 6 | - | 5 | 6 | 6 | 4 |
| $r_{min} (cm)$ | - | - | - | - | - | 4 | 4 | - | 2 | 2 | 2 | 1 |
| $\varphi_0 (deg)$ | - | - | - | - | - | 165 | 165 | - | 140 | 100 | 100 | 80 |
| $\varphi_{max} (deg)$ | - | - | - | - | - | 45 | 45 | - | 45 | 180 | 180 | 180 |
| $\varphi_{min} (deg)$ | 160 | 120 | 150 | 170 | 180 | 125 | 120 | 160 | 165 | 80 | 90 | 110 |
| $\rho (-)$ | 0.5 | 0.5 | 0.5 | 0.3 | 0.5 | 0.5 | 0.6 | 0.7 | 0.7 | 0.7 | 0.5 | 0.7 |

### Excitation-contraction coupling

Excitation-contraction coupling was used to represent physiological activation dynamics of muscle stimulation:

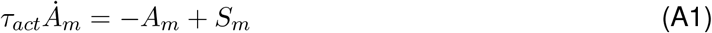

where *τ*_*act*_ = 0.01 when the muscle is activating (*S*_*m*_ *≥* 0) and *τ*_*act*_ = 0.04 when deactivating (*S*_*m*_ *<* 0), set from (Thelen, 2003).

## Appendix B Kinematic Fitting Results

Figure B1 shows the simulated gait obtained from the kinematic fitting optimization used to initialize the nominal walking solution. Simulated kinematics, kinetics, and muscle activations are compared with experimental reference data for walking at 1.2 m s^*−*1^. Compared with the minimum-effort optimization results (Fig. 2), kinematic RMSE was substantially smaller for most control models. In particular, the reflex-based and CPG-reflex-synergy-based models reproduced more human-like early-stance knee flexion, and the reflex-based model no longer exhibited excessive ankle plantarflexion during early swing. However, improvements in kinematic agreement were often accompanied by larger deviations in joint kinetics and muscle activation patterns, suggesting that optimization based primarily on kinematic matching does not necessarily produce more physiologically realistic gait in neuromechanical simulations.

**Figure B1:**
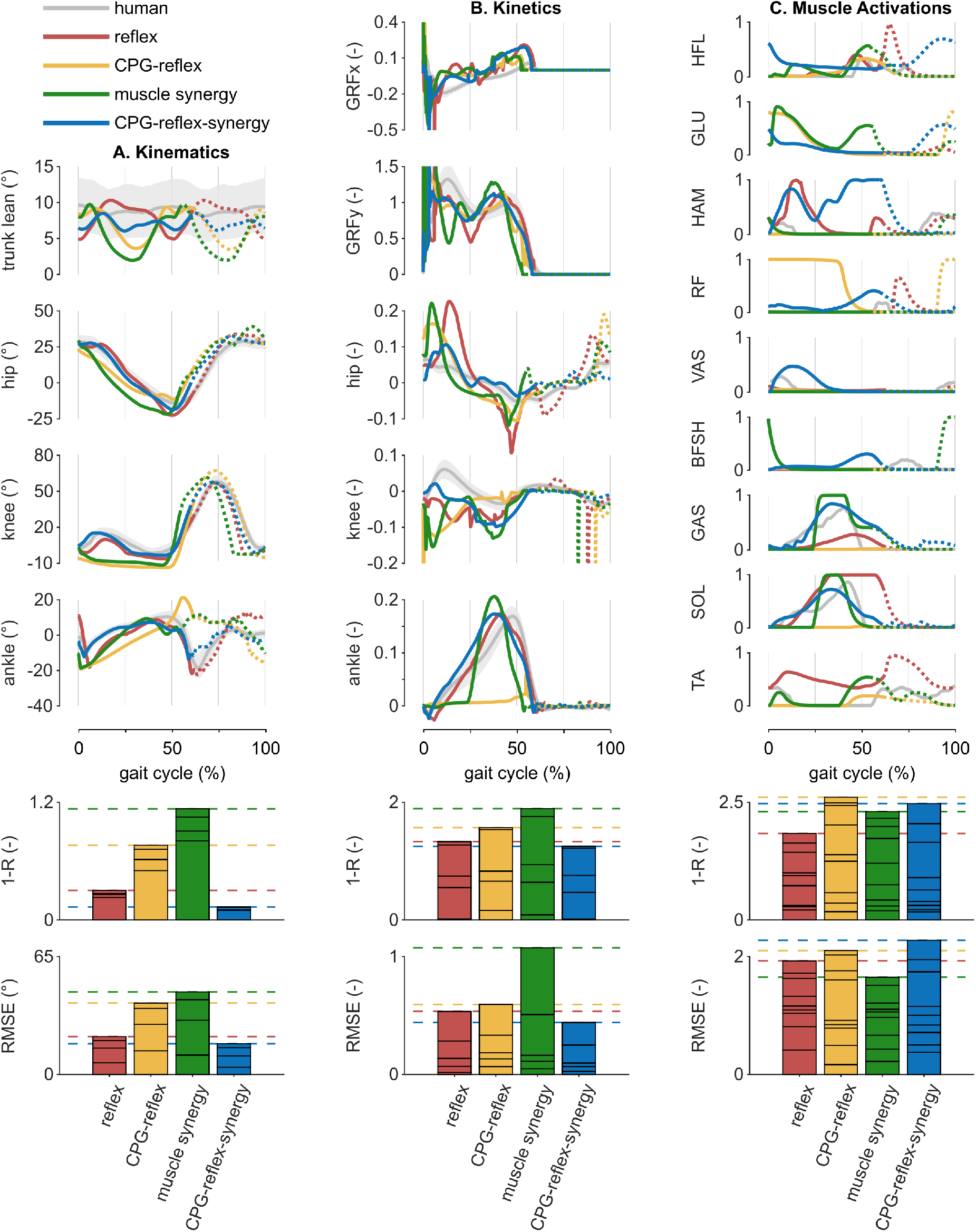
Simulated gait and comparison to reference data for walking at 1.2 m s^*−*1^ obtained from kinematic fitting. Same format as Fig. 2, but showing results from the kinematic fitting optimization used for initialization, whereas Fig. 2 presents results from minimum-effort optimization. Kinematics, kinetics, and muscle activations are compared to experimental reference data, along with corresponding deviations.

## Additional information

### Data availability statement

All data is presented in the figures. The MATLAB/Simulink simulation framework, controller implementations, and optimized parameter sets are available at *https://doi.org/10.5281/zenodo.20509551*, release v1.0.0. An accompanying video demonstration of the simulated locomotion results is available at *https://youtu.be/suMJIfZYe5k?si=GeTNjrnSL4SfAJ-T*

### Author contributions

Both authors designed the research, analyzed the results, drafted the article, and approved the final version of the manuscript. V.T. implemented the computational model and conducted the simulation studies. The authors agree to be accountable for all aspects of the work.

## Acknowledgments

The authors thank A. Di Russo and A. Ijspeert for their guidance on the implementation details of their CPG-reflex-synergy-based model.

## References

Ackermann, M., & Van den Bogert, A. J. (2010). Optimality principles for model-based prediction of human gait. Journal of biomechanics, 43(6), 1055–1060.

Aoi, S., Ogihara, N., Funato, T., Sugimoto, Y., & Tsuchiya, K. (2010). Evaluating functional roles of phase resetting in generation of adaptive human bipedal walking with a physiologically based model of the spinal pattern generator. Biological Cybernetics, 102(5), 373–387.

Aoi, S., Ogihara, N., Sugimoto, Y., & Tsuchiya, K. (2008). Simulating Adaptive Human Bipedal Locomotion Based on Phase Resetting Using Foot-Contact Information. Advanced Robotics, 22(15), 1697–1713.

Aoi, S., Ohashi, T., Bamba, R., Fujiki, S., Tamura, D., Funato, T., Senda, K., Ivanenko, Y., & Tsuchiya, K. (2019). Neuromusculoskeletal model that walks and runs across a speed range with a few motor control parameter changes based on the muscle synergy hypothesis. Scientific Reports, 9(1), 369.

Arnold, E. M., Hamner, S. R., Seth, A., Millard, M., & Delp, S. L. (2013). How muscle fiber lengths and velocities affect muscle force generation as humans walk and run at different speeds. Journal of Experimental Biology.

Bhargava, L. J., Pandy, M. G., & Anderson, F. C. (2004). A phenomenological model for estimating metabolic energy consumption in muscle contraction. Journal of Biomechanics, 37 (1), 81–88.

Browning, R. C., Baker, E. A., Herron, J. A., & Kram, R. (2006). Effects of obesity and sex on the energetic cost and preferred speed of walking. Journal of Applied Physiology, 100(2), 390–398.

Camargo, J., Ramanathan, A., Flanagan, W., & Young, A. (2021). A comprehensive, open-source dataset of lower limb biomechanics in multiple conditions of stairs, ramps, and level-ground ambulation and transitions. Journal of Biomechanics, 119, 110320.

Danner, S. M., Shevtsova, N. A., Frigon, A., & Rybak, I. A. (2017). Computational modeling of spinal circuits controlling limb coordination and gaits in quadrupeds. Elife, 6, e31050.

De Groote, F., & Falisse, A. (2021). Perspective on musculoskeletal modelling and predictive simulations of human movement to assess the neuromechanics of gait. Proceedings of the Royal Society B: Biological Sciences, 288(1946), 20202432.

Di Russo, A., Stanev, D., Sabnis, A., Danner, S. M., Ausborn, J., Armand, S., & Ijspeert, A. (2023). Investigating the roles of reflexes and central pattern generators in the control and modulation of human locomotion using a physiologically plausible neuromechanical model. Journal of Neural Engineering, 20(6), 066006.

Dietz, V. (2002). Proprioception and locomotor disorders. Nature Reviews Neuroscience, 3(10), 781–790.

Dzeladini, F., Van Den Kieboom, J., & Ijspeert, A. (2014). The contribution of a central pattern generator in a reflex-based neuromuscular model. Frontiers in Human Neuroscience, 8.

Feldman, A. G., & Levin, M. F. (1995). The origin and use of positional frames of reference in motor control. Behavioral and Brain Sciences, 18(4), 723–744.

Fleischmann, S., Shanbhag, J., Miehling, J., Wartzack, S., Ong, C., Eskofier, B. M., & Koelewijn, A. D. (2025). A model of the cerebellum generates gait adaptations in a reflex-based neuromusculoskeletal model during split-belt walking. Journal of NeuroEngineering and Rehabilitation, 22(1), 256.

Franz, J. R., Lyddon, N. E., & Kram, R. (2012). Mechanical work performed by the individual legs during uphill and downhill walking. Journal of Biomechanics, 45(2), 257–262.

Fritz, S., & Lusardi, M. (2009). White paper: “walking speed: The sixth vital sign”. Journal of Geriatric Physical Therapy (2001), 32(2), 46–49.

Geijtenbeek, T. (2019). SCONE: Open Source Software for Predictive Simulation of Biological Motion. Journal of Open Source Software, 4(38), 1421.

Geyer, H., & Herr, H. (2010). A Muscle-Reflex Model That Encodes Principles of Legged Mechanics Produces Human Walking Dynamics and Muscle Activities. IEEE Transactions on Neural Systems and Rehabilitation Engineering, 18(3), 263–273.

Grillner, S. (2006). Biological Pattern Generation: The Cellular and Computational Logic of Networks in Motion. Neuron, 52(5), 751–766.

Günther, M., & Ruder, H. (2003). Synthesis of two-dimensional human walking: A test of the λ − model. Biological Cybernetics, 89(2), 89–106.

Haeufle, D. F., Schmortte, B., Geyer, H., Müller, R., & Schmitt, S. (2018). The benefit of combining neuronal feedback and feed-forward control for robustness in step down perturbations of simulated human walking depends on the muscle function. Frontiers in computational neuroscience, 12, 80.

Hansen, N. (2006). The CMA Evolution Strategy: A Comparing Review [Series Title: Studies in Fuzziness and Soft Computing]. In J. A. Lozano, P. Larrañaga, I. Inza, & E. Bengoetxea (Eds.), Towards a New Evolutionary Computation (pp. 75–102, Vol. 192). Springer Berlin Heidelberg.

Hase, K., & Yamazaki, N. (1997). Development of Three-Dimensional Whole-Body Musculoskeletal Model for Various Motion Analyses. JSME international journal. Ser. C, Dynamics, control, robotics, design and manufacturing, 40(1), 25–32.

Hase, K., & Yamazaki, N. (2002). Computer simulation study of human locomotion with a three-dimensional entire-body neuro-musculo-skeletal model (I. Acquisition of normal walking). JSME International Journal Series C Mechanical Systems, Machine Elements and Manufacturing, 45(4), 1040–1050.

Hirasaki, E., Moore, S. T., Raphan, T., & Cohen, B. (1999). Effects of walking velocity on vertical head and body movements during locomotion. Experimental Brain Research, 127 (2), 117–130.

Ijspeert, A. J. (2008). Central pattern generators for locomotion control in animals and robots: A review. Neural Networks, 21(4), 642–653.

Ivanenko, Y. P., Poppele, R. E., & Lacquaniti, F. (2004). Five basic muscle activation patterns account for muscle activity during human locomotion. The Journal of Physiology, 556(1), 267–282.

Jo, S., & Massaquoi, S. G. (2007). A model of cerebrocerebello-spinomuscular interaction in the sagittal control of human walking. Biological Cybernetics, 96(3), 279–307.

Jones, R., Ratnakumar, N., Akbaş, K., & Zhou, X. (2024). Delayed center of mass feedback in elderly humans leads to greater muscle co-contraction and altered balance strategy under perturbed balance: A predictive musculoskeletal simulation study (A. Mengarelli, Ed.). PLOS ONE, 19(5), e0296548.

Kim, Y., Tagawa, Y., Obinata, G., & Hase, K. (2011). Robust control of CPG-based 3D neuromus-culoskeletal walking model. Biological Cybernetics, 105(3-4), 269–282.

Koelewijn, A. D., Heinrich, D., & van den Bogert, A. J. (2019). Metabolic cost calculations of gait using musculoskeletal energy models, a comparison study. PloS one, 14(9), e0222037.

Lankford, D. E., Wu, Y., Bartschi, J. T., Hathaway, J., & Gidley, A. D. (2020). Development and validation of a steep incline and decline metabolic cost equation for steady-state walking. European Journal of Applied Physiology, 120(9), 2095–2104.

Matsuoka, K. (1985). Sustained oscillations generated by mutually inhibiting neurons with adaptation. Biological Cybernetics, 52(6), 367–376.

Miller, R. H. (2014). A comparison of muscle energy models for simulating human walking in three dimensions. Journal of Biomechanics, 47 (6), 1373–1381.

Nielsen, J. B. (2003). How we Walk: Central Control of Muscle Activity during Human Walking. The Neuroscientist, 9(3), 195–204.

Ogihara, N., & Yamazaki, N. (2001). Generation of human bipedal locomotion by a bio-mimetic neuro-musculo-skeletal model. Biological Cybernetics, 84(1), 1–11.

Ong, C. F., Geijtenbeek, T., Hicks, J. L., & Delp, S. L. (2019). Predicting gait adaptations due to ankle plantarflexor muscle weakness and contracture using physics-based musculoskeletal simulations (M. Srinivasan, Ed.). PLOS Computational Biology, 15(10), e1006993.

Paul, C., Bellotti, M., Jezernik, S., & Curt, A. (2005). Development of a human neuro-musculo-skeletal model for investigation of spinal cord injury. Biological Cybernetics, 93(3), 153–170.

Perry, J., & Burnfield, J. M. (2010). Gait analysis: Normal and pathological function (2nd ed).

SLACK.

Ramadan, R., Geyer, H., Jeka, J., Schöner, G., & Reimann, H. (2022). A neuromuscular model of human locomotion combines spinal reflex circuits with voluntary movements. Scientific Reports, 12(1), 8189.

Rose, J., & Gamble, J. G. (Eds.). (2006). Human walking (3rd ed). Lippincott Williams & Wilkins.

Severini, G., & Muñoz, D. (2025). A physiologically inspired hybrid CPG/Reflex controller for cycling simulations that generalizes to walking (G. Durandau, Ed.). PLOS Computational Biology, 21(9), e1013494.

Song, S., & Geyer, H. (2012). Regulating speed and generating large speed transitions in a neuromuscular human walking model. 2012 IEEE International Conference on Robotics and Automation, 511–516.

Song, S., & Geyer, H. (2015a). A neural circuitry that emphasizes spinal feedback generates diverse behaviours of human locomotion. The Journal of Physiology, 593(16), 3493–3511.

Song, S., & Geyer, H. (2015b). Regulating speed in a neuromuscular human running model. 2015 IEEE-RAS 15th International Conference on Humanoid Robots (Humanoids), 217–222.

Song, S., & Geyer, H. (2017). Evaluation of a Neuromechanical Walking Control Model Using Disturbance Experiments. Frontiers in Computational Neuroscience, 11.

Song, S., & Geyer, H. (2018). Predictive neuromechanical simulations indicate why walking performance declines with ageing. The Journal of Physiology, 596(7), 1199–1210.

Song, S., Kidziński, Ł., Peng, X. B., Ong, C., Hicks, J., Levine, S., Atkeson, C. G., & Delp, S. L. (2021). Deep reinforcement learning for modeling human locomotion control in neurome-chanical simulation. Journal of NeuroEngineering and Rehabilitation, 18(1), 126.

Taga, G. (1995). A model of the neuro-musculo-skeletal system for human locomotion: I. Emergence of basic gait. Biological Cybernetics, 73(2), 97–111.

Tamura, D., Aoi, S., Funato, T., Fujiki, S., Senda, K., & Tsuchiya, K. (2020). Contribution of Phase Resetting to Adaptive Rhythm Control in Human Walking Based on the Phase Response Curves of a Neuromusculoskeletal Model. Frontiers in Neuroscience, 14, 17.

Thelen, D. G. (2003). Adjustment of Muscle Mechanics Model Parameters to Simulate Dynamic Contractions in Older Adults. Journal of Biomechanical Engineering, 125(1), 70–77.

Ton, V., Solav, D., & Song, S. (2024). Impact of Sole Designs of Offloading AFO on Gait Dynamics: A Predictive Neuromechanical Simulation Study. 2024 10th IEEE RAS/EMBS International Conference for Biomedical Robotics and Biomechatronics (BioRob), 1709–1714.

Tresch, M. C., & Jarc, A. (2009). The case for and against muscle synergies. Current Opinion in Neurobiology, 19(6), 601–607.

Uchida, T. K., Seth, A., Pouya, S., Dembia, C. L., Hicks, J. L., & Delp, S. L. (2016). Simulating Ideal Assistive Devices to Reduce the Metabolic Cost of Running (Ø. Sandbakk, Ed.). PLOS ONE, 11(9), e0163417.

Umberger, B. R. (2010). Stance and swing phase costs in human walking. Journal of The Royal Society Interface, 7 (50), 1329–1340.

Umberger, B. R., Gerritsen, K. G., & Martin, P. E. (2003). A Model of Human Muscle Energy Expenditure. Computer Methods in Biomechanics and Biomedical Engineering, 6(2), 99–111.

Veerkamp, K., Waterval, N., Geijtenbeek, T., Carty, C., Lloyd, D., Harlaar, J., & Van Der Krogt, M. (2021). Evaluating cost function criteria in predicting healthy gait. Journal of Biomechanics, 123, 110530.

Wang, J. M., Hamner, S. R., Delp, S. L., & Koltun, V. (2012). Optimizing locomotion controllers using biologically-based actuators and objectives. ACM Transactions on Graphics, 31(4), 1–11.

Waterval, N., Brehm, M., Veerkamp, K., Geijtenbeek, T., Harlaar, J., Nollet, F., & Van Der Krogt, M. (2023). Interacting effects of AFO stiffness, neutral angle and footplate stiffness on gait in case of plantarflexor weakness: A predictive simulation study. Journal of Biomechanics, 157, 111730.

Winter, D. A. (2009). Biomechanics and motor control of human movement (4th ed). Wiley.

Yamazaki, N., Hase, K., Ogihara, N., & Hayamizu, N. (1996). Biomechanical analysis of the development of human bipedal walking by a neuro-musculo-skeletal model. Folia Primatologica, 66(1-4), 253–271.

